# Ontogeny of settlement behaviours in response to *Grammatophora marina* diatom biofilms in the marine polychaete, *Platynereis dumerilii*

**DOI:** 10.64898/2026.04.10.717688

**Authors:** Callum Teeling, Susanne Vogeler, Robert P Ellis, Elizabeth A Williams

## Abstract

Settlement, the transition of a swimming planktonic larva to a crawling or sessile benthic juvenile, is a key process in the development of many marine invertebrates. Successful recruitment via larval settlement is critical for the development and maintenance of seafloor ecosystems. Microbial biofilms act as positive cues for larval settlement across diverse taxa, yet the behavioural processes preceding settlement are poorly understood. Here, we investigated age-dependent changes in settlement behaviour in the marine polychaete *Platynereis dumerilii* larvae in response to *Grammatophora marina* diatom biofilms. Settlement behaviours (crawling, crawling speed, and track straightness (tortuosity)) were quantified from recordings of larvae at five developmental stages (mid-trochophore to late-nectochaete) in the presence or absence of diatom biofilms, using image segmentation and spot-tracking software. As larvae developed, the proportion of individuals crawling (settlement) over the biofilm increased. Older larvae colonised biofilms more rapidly and showed greater discrimination between *G. marina* biofilms and non-biofilmed controls. The movement trajectory of older larvae also straightens compared to individuals swimming in the presence of biofilms, or behaviour witnessed in the absence of biofilms. The proportions and magnitudes of these behaviours may reflect changing prioritisation of sensory inputs from physical and chemical cues as larvae develop. Our findings suggest that behavioural traits that are associated with settlement are developmentally programmed in *P. dumerilii*. Understanding settlement behaviours in *P. dumerilii* expands on this species’ behavioural repertoire and sheds light on the evolutionary relationship between marine larvae and microalgal biofilms.

## Introduction

Settlement is a crucial transition in the life cycle of many marine invertebrates. It is defined as a reversible behaviour which involves an individual switching from swimming in the plankton to crawling on and exploring the seafloor, followed by the permanent metamorphosis into a benthic juvenile (Crisp, 1974). Settlement occurs once larvae reach competency - the developmental stage at which larvae descent from the water column and are physiologically capable of completing metamorphosis. The ecology and health of many marine ecosystems are dictated by settlement success. This success depends on a variety of factors such as finding adequate settlement sites, outcompeting other individuals for space, and finding sufficient nutrients to survive metamorphosis into the juvenile life stage (Chong-Seng et al., 2014; Gosselin & Qian, 1996; Méndez Casariego et al., 2004; Toone et al., 2023). Settlement is believed to be driven by environmental cues such as chemical signals produced by conspecifics, macroalgae, or microbial biofilms (Hadfield & Paul, 2001; Pawlik, 1992). However, a deeper understanding of which cues induce settlement and how those cues affect settlement decisions is still missing.

One of the key biological settlement cues in the oceans are microbial biofilms. These can form on any surface or substrate, and are often associated with rocky shores, reefs, mud and sand, as well as forming on other biota such as macroalgae (Pawlik, 1992). Microbial biofilms are first established by bacteria which are then quickly colonised by diatoms which utilise the extracellular polymeric substances from the bacteria as an attachment substrate and nutrient source (Antunes et al., 2019; Zobell & Allen, 1935). This succession can change the biofilm composition, structure and surface properties, altering the components as potential settlement inducers for other species. Despite their contribution to benthic habitats, the role of microalgal biofilms in marine larval settlement and metamorphosis have historically received less attention than microbial equivalents. Studies exposing marine invertebrate larvae to single species, or mixed, diatom biofilms show an increase in larval settlement in the annelid *Hydroides elegans* (Harder et al., 2002; Lam et al., 2005), the bryozoan *Bugula neritina* (Dahms et al., 2004; Kitamura & Hirayama, 1987), and several mollusc species (Daume et al., 2000; Takami et al., 1997; Wang et al., 2012).

Moreover, research on the effects of specific diatom biofilms for larval settlement has aided in the commercial production of economically important invertebrate species. For example, collection nets coated in the diatoms *Fragilariopsis pseudonana* and *Navicula veneta* increase the number of *Argopecten purpuratus* scallop spat collected (Avendaño-Herrera et al., 2003). Despite evidence of the importance of diatom biofilms for inducing settlement, questions remain regarding how the biofilms trigger settlement and surface exploration behaviours. With many experiments having focused on settlement rates of larvae following 24 h of exposure to microalgal biofilms (Daume et al., 2000; Freckelton et al., 2024; Lau et al., 2005; Unabia & Hadfield, 1999), it is possible we may be missing key behavioural traits displayed by larvae that precede their final settled state.

Some of the best described surface exploration behaviours in a marine larva exists for barnacle cyprids (Aldred et al., 2018; Crisp, 1976; Maleschlijski et al., 2012, 2015). Barnacle cyprids will often swim with high velocity in straight trajectories or spirals, interspersed with sinking behaviours, to initiate surface contact. Once contact with a surface cue is made, cyprids will then explore the surface by walking on their antennules with low velocity or crawling over the surface while rotating with a medium velocity. Biofilm exploration and settlement behaviour has also been described in the polychaete, *H. elegans*. Bacterial biofilms are shown to increase the number of larvae crawling over and touching the surface, with crawling characterised as tracks with lower velocity and fewer turns compared to swimming tracks (Hadfield et al., 2014). Despite these efforts, information is still lacking on biofilm exploration behaviours across different marine invertebrate larvae. Even less knowledge is available for how these behaviours might change over the course of larval development.

The marine polychaete, *Platynereis dumerilii*, is a well-established laboratory model organism which has been used to study behaviour, neurobiology, development, and evolution (Özpolat et al., 2021). *P. dumerilii* belongs to the sub-phylum of Lophotrochozoa which also includes molluscs, brachiopods, bryozoans, and nemerteans. Like many lophotrochozoans, *P. dumerilii* develops through free-swimming and fully pelagic trochophore stages which show no crawling behaviour. By the early to mid-nectochaete stages parapodia and crawling muscles have developed, and by late-nectochaete the larvae live mostly on the seafloor, at which point feeding is initiated (Fischer et al., 2010). As interest in the sensory biology of marine ciliated larvae increases, *P. dumerilii* is emerging as a powerful laboratory model in this field (e.g. Bezares Calderón et al., 2024; Bezares-Calderón et al., 2018; Chartier et al., 2018; Gühmann et al., 2015; Jokura et al., 2023).

Recently, the settlement preferences of *P. dumerilii* larvae have been established, showing *P. dumerilii* to be a generalist, settling on a range of diatom biofilms, but with a species-specific response to different diatoms. The chain-forming diatom, *Grammatophora marina*, induces the highest median levels of settlement after 24 h of exposure (Hird et al., 2024). *Grammatophora spp.* have been identified as abundant epiphytes on the seagrass, *Posidonia oceanica*, within the Bay of Naples, a habitat where *P. dumerilii* reside, living on the seagrass blades (Gambi et al., 2000; Mutalipassi et al., 2020). *Grammatophora spp.* diatoms have also been identified in the faecal pellets of laboratory reared adults (Gambi et al., 2000). A recent study shows that *P. dumerilii* larvae exposed to *G. marina* biofilms from an early age can allocate more resources towards nervous system development and growth compared to larvae not exposed to *G. marina* (Randel, 2025). *G. marina* biofilms are therefore likely to be both an ecologically relevant larval settlement cue and juvenile/adult food source for *P. dumerilii*. However, much like for other invertebrates, knowledge on how larvae identify and interact with their preferred settlement cue, prior to committed settlement and attachment, is still lacking.

Here, we used video recording, image segmentation and automated tracking software to characterise specific aspects of settlement behaviour in *P. dumerilii* larvae at different stages of development by exposing them to *G. marina* biofilms. Several settlement behaviours could then be identified from the larval tracks including swimming and crawling speeds, the proportion of larvae crawling across the biofilm, and the straightness (tortuosity) of swimming and crawling tracks. We also assessed larval choice between *G. marina* biofilms and controls across different stages of development. We aimed to characterise biofilm exploration in this species and how these behaviours change with age, with the analysis pipeline developed here transferable to the study of settlement behaviours in other marine larvae.

## Materials and Methods

### *Platynereis dumerilii* culture

*Platynereis dumerilii* were cultured at the Marine Invertebrate Culture Unit (MICU) at the University of Exeter following an adapted protocol (Fischer & Dorresteijn, 2004; Hird et al., 2024). Premature adult atoke and mature adult epitoke worms were housed in 4 L styrene-acrylonitrile boxes (Vitlab) with 1.5 L of UV sterilised, 0.2 µm filtered artificial seawater (FASW, Tropic Marin Pro-Reef salt) at 33 ± 1 ppt and 22 ± 0.5 °C. The adult cultures had a light regime of 16h:8h L:D with a rotating 7-day moon cycle of 30% illumination every three weeks. Worms were fed twice a week on 10 mL of mixed microalgae (*Tetraselmis suecica*, *Nannochloropsis salina*, and *Phaeodactylum tricornutum*), 10 mL of rotifers, or 5 mL rotifer and spinach mix depending on the worm feeding stage. Larval cultures were produced by spawning one female and one male epitoke in FASW until eggs and sperm were released. After fertilisation, eggs were rinsed in FASW to remove excess sperm and incubated at 18 °C. After 24 h, the egg jelly was removed, and successfully hatched larvae were kept in FASW at 18 °C with 16h:8h L:D.

### *Grammatophora marina* culture

The *G. marina* stock culture was obtained from the Culture Collection of Algae and Protozoa (CCAP 1027/1), Scotland. *G. marina* was cultured in 50 mL vented flasks (BioLite) with 40 mL of culture media made from 0.5 L of autoclaved, 0.2 µm filtered artificial seawater (FASW) with 1 mL of Cell-Hi-F2P media (Varicon Aqua, UK) and 200 µL of ZM Silicate Solution (ZM Systems, UK). Flasks were kept in an incubator at 18°C with fluorescent illumination under a 16:8h L:D cycle. Diatoms were sub-cultured every three weeks by transferring 10 mL of culture into a fresh flask, then topping up to 40 mL with fresh F/2 + silica culture media.

### Preparation of *G. marina* coated slips

Six mm discs were cut from sheets of ACLAR film (Agar Scientific, AGL4458). ACLAR slips were autoclaved, then incubated at room temperature for 1 h in 5 µg mL^−1^ poly-D-lysine (Sigma-Aldrich, P6407), rinsed three times in autoclaved MilliQ water with gentle swirling, blotted, and left to dry completely at room temperature for 2 h. ACLAR slips were kept at 4°C for up to 1 month before inoculating with *G. marina*.

To prepare the biofilms, 1.5 mL of *G. marina* cell suspension were pipetted into 24-well plates (Sarstedt, 83.3922.500), each well containing two ACLAR slips. Negative control slips were prepared in 1.5 mL of *G. marina* culture media (F/2 media + silica), in the absence of diatoms. Plates were incubated for 24 h at 18°C. Prior to an experiment, both diatom coated and negative control slips (‘F/2 control’) were carefully rinsed by dipping the slips three times in 0.2 µm FASW to remove loosely adhered cells or precipitated silica. To later determine the coverage of diatoms on the slips, photographs of each slip were taken using an Axiocam 208 Color (Ziess, Germany) with Labscope software, attached to a Stemi 508 stereo microscope (Ziess, Germany). The cell coverage percentage on each slip was calculated using “Biofilm_coverage_macro.ijm” (Hird et al., 2024).

### Recording stage design

Larval behaviours in response to both biofilm coated and control slips were recorded using a custom behavioural set-up (Fig. 1A). The recording stage was built upon an aluminium breadboard using various optomechanical components and supports (Thorlabs, UK). The top horizontal support held a U3-3680XCP-NIR-GL infrared camera with an 8 mm lens (IDS, Germany) and a longpass filter (λ=715 nm, Thorlabs). The lower horizontal support held a small stage which was positioned 8 cm away from the camera. A single infrared ring light (λ=850 nm) with a brightness control adaptor (AmScope, UK) was set beneath the stage.

**Figure 1.**
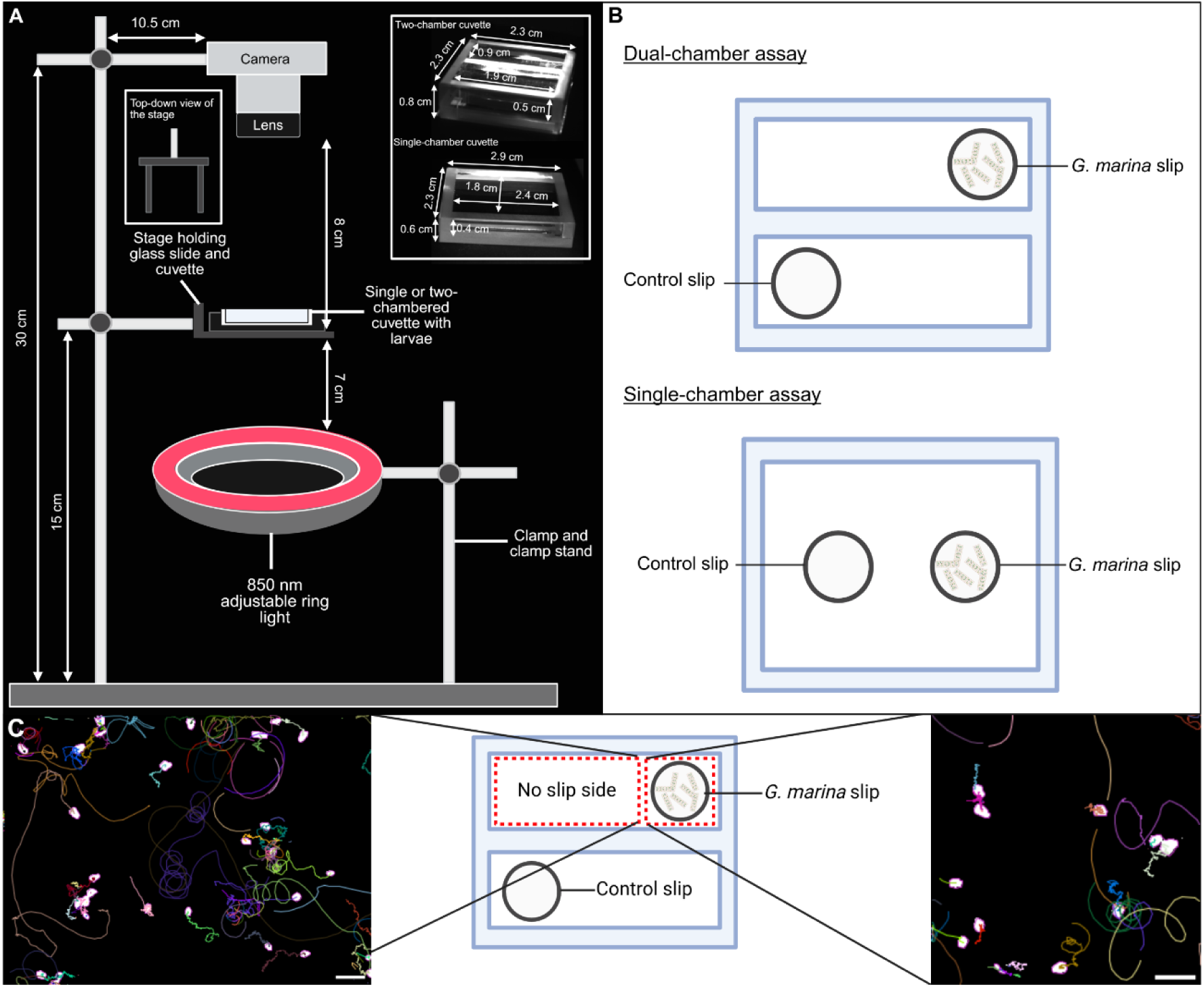
Schematics of the larval recording stage and optical cuvettes used in the experiments. (A) Configuration of the recording stage with measurements with inserts of the optical cuvettes and cuvette holder. (B) Schematic comparison of the dual-chamber and single-chamber cuvettes with example slip placement. (C) The dual-chamber cuvette showing the two ROIs (no-slip and slip) with larval tracks exported from TrackMate showing an example 30-second video clip of 6-dpf larvae, track colours are random and the scale bars = 1 mm. Created with BioRender.com.

### *P. dumerilii* larval behaviour assays

Larval behaviour assays were carried out in bespoke optical cuvettes (Thorlabs, Fig 1B). The dual-chamber assay used a cuvette with two chambers that were each 1.9 x 0.9 x 0.5 cm (L x W x H). A single diatom coated or control ACLAR slip was placed at the far end of one of the chambers. Which chamber the diatom coated and F/2 control slip were placed in (top or bottom) and their position within the chamber (left or right side) was determined by a random number generator. Each chamber was filled with 900 µL of 0.2 µm FASW and between 30-60 *P. dumerilii* were pipetted into each chamber at the opposite end from the slip.

The single-chamber assay used an optical cuvette with only a single chamber with dimensions 2.4 x 1.8 x 0.4 cm. In this assay, both the diatom coated and control slips were set at opposite ends of the cuvette with the position of each slip again determined by a random number generator. The chamber was filled with 1.8 mL of 0.2 µm FASW and between 50 – 150 larvae were pipetted into the centre of the chamber. For both assays, larvae were given a 2 min acclimation period to recover from physical handling and pipetting disturbances before recording commenced.

Both assays were repeated six times, with each replicate made using age-matched larvae pooled from at least two different batches with different parentage. This equalled twelve larval batches per age for all replicates (72 batches total across all experiments). Experiments were conducted with 2-days post-fertilisation (dpf) trochophore larvae, 3-, 3.5-, 4-dpf mid-nectochaete larvae, and 5-, 6-dpf late-nectochaete larvae to test whether settlement behaviours changed with age. These ages were selected based on Hird et al. (2024) who showed that settlement begins between 3- and 4-dpf, and to test if 2-dpf larvae change their behaviour in response to diatoms even if they do not settle upon the biofilm. The six replicate experiments for each age group were performed on different days. The cuvettes were placed onto the horizontal stage on top of a glass slide that was taped under black card. A 2.5 x 2.5 cm square hole was cut within the card that each cuvette could sit within. Larvae were recorded for 5 minutes at 20 frames per second using IDS Peak Cockpit software v1.6.0.0. Videos were made under infrared illumination and in near darkness to avoid any effect of phototaxis in younger larvae (Fischer et al., 2010).

### Data analysis

Videos were prepared for analysis in FIJI v2.14.0/1.54i. For the dual-chamber assay, each chamber and area was analysed separately (Fig. 1C). The slip and no-slip regions of interest (ROIs) were selected using the rectangle selection tool. The background was subtracted from the ROIs, and the larvae were thresholded using an ImageJ macro (Create_Mask.ijm). Each larva was now represented as a white spot against a black background. Video analysis of larval behaviour was performed in FIJI with the TrackMate 7 plugin (Ershov et al., 2022) using the mask detector and simple LAP tracker with default settings. Track details for each larva were exported and further analysed.

Before analysis, tracks that had a duration of less than 20 frames (1 sec of video) were removed. After initial manual inspection of the track speed data against the video data, it was determined that the crawling speed of *P. dumerilii* larvae did not exceed a mean of 0.24 mm/s at any age tested. The proportion of crawling tracks over each area was calculated using the threshold of ≤ 0.24 mm/s with the proportion of all swimming tracks calculated with the threshold of > 0.24 mm/s. Plots of the average track speed data can be seen in Supplementary Figure 1. To assess track straightness, we calculated the average track tortuosity (Codling et al., 2008) using modified R scripts (Bezares Calderón et al. 2024).

We also determined if the number of larvae on the diatom coated slips and F/2 control slips changed over time. To do this, we calculated the number of spots per frame normalised to the peak spot number of an entire 5-minute video. We called this measurement the Relative Settled State or RSS (Lilly et al., 2024). Linear models were fitted to the raw data, and the slopes were calculated for each fitted line.

For the single-chamber assay, the number of larvae on each slip, and the number of larvae not on a slip at all, were counted after 5-minutes of video. The number of larvae on the slips and not on slips was then converted to a percentage of the total number of larvae in the cuvette.

### Statistical analysis

All statistical analyses were carried out using RStudio v.4.3.3 (R Core Team, 2024). We fitted generalised linear models (GLM) using beta distribution and a logit link function to analyse the effect of age, treatment, and area on the proportion of crawling larvae and track tortuosity. A gaussian GLM was used to analyse the same fixed effects on larval crawling speeds. We also fitted a beta GLM with logit link function to test the interactive effects of age and treatment on RSS. Model performances were assessed using Akaike Information Criterion (AIC) and goodness-of-fit was evaluated from the residual deviance. The final models used for analysis and their outputs are shown in Supplementary Tables 1-4. Pairwise comparisons between model variables were performed using the emmeans package (Lenth & Piaskowski, 2025). The effects of age and treatment on the tortuosity of swimming and crawling tracks were determined by one-way ANOVA after confirmation that model assumptions were met. Pairwise comparisons between ages and treatments were carried out using Tukey’s HSD post-hoc analysis. To test if diatom coverage influences crawling proportions, crawling speeds, and tortuosity, linear regression models were fitted for all behaviours at each age.

Choice preference data were arcsine square root transformed to satisfy the assumption of normality. Pairwise comparisons between ages and slips were carried out using Tukey’s HSD post-hoc analysis. All TrackMate analysis pipelines and data analysis scripts are available on GitHub (https://github.com/MolMarSys-Lab/Platynereis-settlement_Teeling_et_al_2026).

## Results

Larval movements were successfully tracked with our custom set-up using an infrared camera and the Trackmate 7 plugin in Fiji. Our video analysis pipeline can distinguish between swimming and crawling *P. dumerilii* larvae across multiple ages, including their movement speeds and tortuosity on biofilm and negative control slips and in no-slip areas.

Before interrogating changes in larval behaviours across the different ROIs and treatments from the dual-chamber assay, we wanted to understand the level of baseline behaviour in *P. dumerilii* larvae. To do this, we analysed tracks from the no-slip side of the chamber containing a negative control slip (‘F/2 control’) in the dual-chamber assay (Fig. 1C). We hypothesised that this ROI would be indicative of larval behaviour in the absence of any biofilm associated cues or physical, textural cues from the control ACLAR slip. Descriptions of the maximum and minimum mean track speeds, proportion of swimming and crawling larvae based on a 0.24 mm/s threshold, and the mean tortuosity of larval tracks can be seen in Table 1. Overall, swimming speeds and the proportion of crawling larvae increased with age, while track tortuosity decreased with age.

**Table 1.** Descriptions of swimming and crawling behaviours, and track tortuosity (straightness) of *P. dumerilii* larvae from 2- to 6-days post-fertilisation (dpf) in the absence of *G. marina* biofilm and ACLAR slip. * 2-dpf larvae cannot crawl so this speed is not indicative of crawling based on our 0.24 mm/s threshold for older larvae.

| Larval age (dpf) | Maximum mean track speed (mm/s) | Minimum mean track speed (mm/s) | Proportion of swimming larvae (%) | Proportion of crawling larvae (%) | Mean track tortuosity |
| --- | --- | --- | --- | --- | --- |
| 2 | 2.700 | 0.011* | 100 | N/A | 0.807 |
| 3 | 2.533 | 0.014 | 88 | 12 | 0.809 |
| 3.5 | 2.974 | 0.001 | 81 | 19 | 0.751 |
| 4 | 3.125 | 0.003 | 76 | 24 | 0.596 |
| 5 | 3.067 | 0.003 | 81 | 19 | 0.629 |
| 6 | 2.824 | 0.003 | 78 | 22 | 0.629 |

### Effect of diatom biofilm and larval age on behaviour

Results from the GLM revealed that there was an interactive effect of larval age and the presence of *G. marina* biofilm on the proportion of crawling larvae. The impact of the biofilm presence on the proportion of crawling larvae becomes increasingly positive with increasing larval age (Fig. 2A, Supplementary Table 1). This relationship becomes significant at 4-dpf (estimate = 1.258 ± 0.396, z = 3.181, *p* < 0.01), with 5-dpf (estimate = 1.423 ± 0.397, z = 3.582, *p* < 0.001) and 6-dpf (estimate = 1.215 ± 0.396, z = 3.070, *p* < 0.01) also significantly different compared to 3-dpf larvae in the F/2 control chamber. The model also shows that the slip had a strong additive effect on the proportion of crawling larvae compared to the no-slip area for both F/2 control and *G. marina* treatments (estimate = 0.913 ± 0.124, z = 7.385, *p* < 0.001). Pairwise comparisons revealed that there were significantly fewer crawling larvae in the F/2 control chamber compared to the *G. marina* chamber at 4-dpf (estimate = -0.901 ± 0.269, z = -3.345, *p* < 0.01), 5-dpf (estimate = -1.066 ± 0.272, z = -3.922, *p* < 0.001), and 6-dpf. Within the F/2 control chamber there were no differences in crawling larvae between ages. In the *G. marina* chamber, there were no significant differences between 4-, 5-, and 6-dpf.

**Figure 2.**
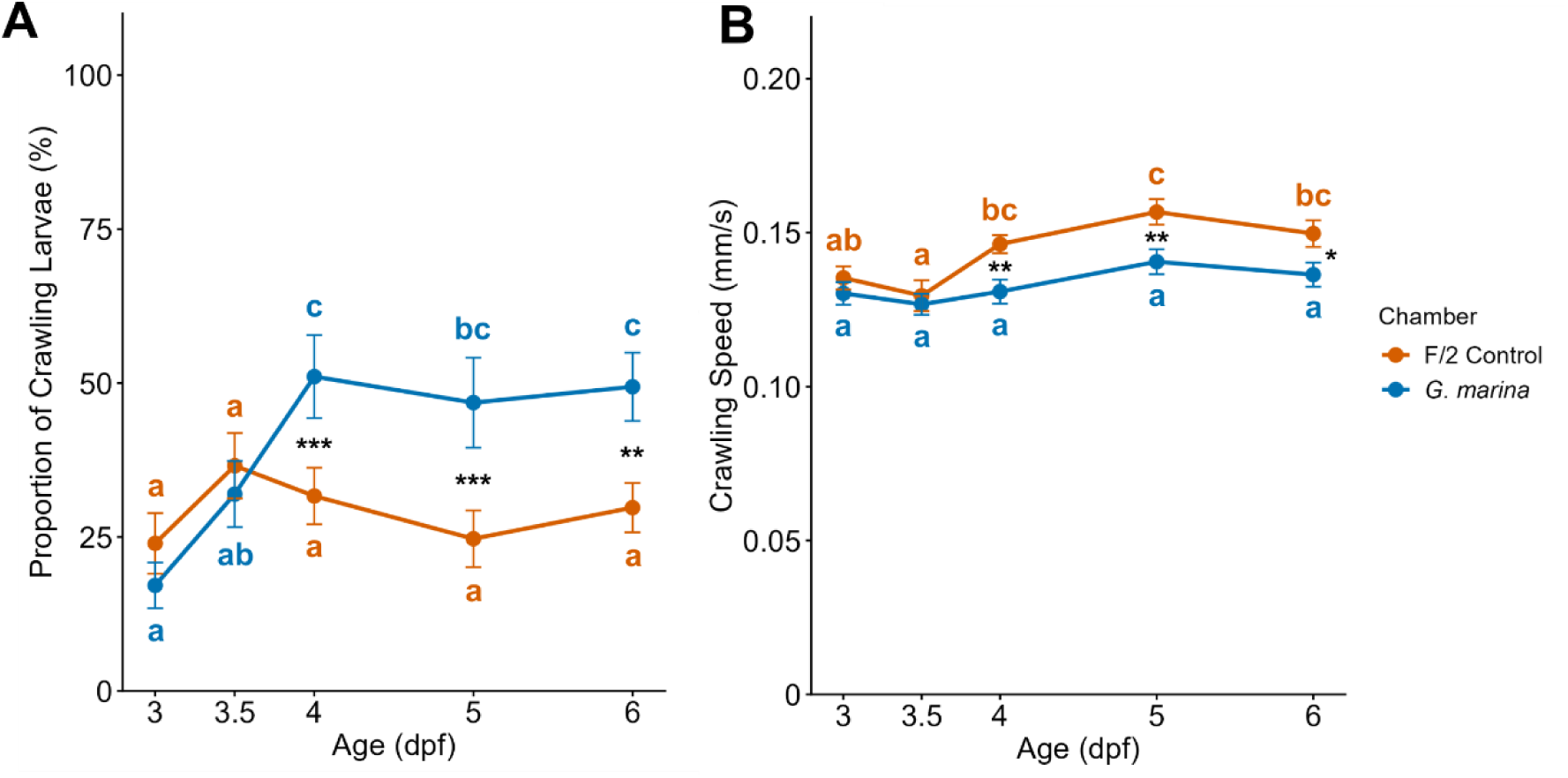
The effect of the *G. marina* biofilm on larval crawling behaviour by age. (A) The proportion of crawling larvae and (B) the crawling speed of larvae within the F/2 control chamber (orange) and the *G. marina* chamber (blue). The ages of *P. dumerilii* larvae are expressed as days post-fertilisation (dpf). Data for 2-dpf larvae was removed as larvae at this age do not crawl. Different letters of the same colour represent statistically significant differences within chambers across ages. Asterisks indicate statistically significant differences between treatments within ages where * *p* < 0.05, ** *p* < 0.01, *** *p* < 0.001. *N* = 6 videos for each age group.

The GLM analysis for crawling speed revealed that the interactive effect between age and *G. marina* biofilm reduced the crawling speed of larvae at 3.5-, 4-, 5-, and 6-dpf (Fig. 2B, Supplementary Table 2). However, this interaction was not significant compared to 3-dpf larvae in the F/2 control chamber. Age had the strongest effect on crawling speed with larvae aged 4-dpf (estimate = 0.011 ± 0.005, t_109_ = 2.055, *p* < 0.05), 5-dpf (estimate = 0.021 ± 0.005, t_109_ = 4.016, *p* < 0.001), and 6-dpf (estimate = 0.144 ± 0.005, t_109_ = 2.702, *p* < 0.01) having significantly higher crawling speeds compared to 3-dpf larvae. The slip area also had a significant, negative effect on crawling speed compared to the no-slip area of the chambers across all ages and treatments (estimate = -0.008 ± 0.002, t_109_ = -3.432, *p* < 0.001). Despite this, pairwise comparisons revealed that crawling speeds in the control chamber were significantly faster compared to the *G. marina* chamber for ages 4-dpf (estimate = 0.015 ± 0.005, t_109_ = 2.894, *p* < 0.01), 5-dpf (estimate = 0.016 ± 0.005, t_109_ = 3.034, *p* < 0.01), and 6-dpf (estimate = 0.013 ± 2.506, *p* < 0.05). In the F/2 control chamber, there was a significant increase in crawling speed from 4-dpf onwards. In the *G. marina* chamber, crawling speeds did not differ between any age of larvae.

A three-way interaction between age, treatment and area was investigated by GLM on larval track tortuosity. Here, the model revealed no significant interactions between age, *G. marina* biofilm, and the slip area on tortuosity when compared to 2-dpf larvae in the F/2 control chamber and no-slip area (Supplementary Table 3). However, there was a strong effect of age which showed reduced track tortuosity, i.e. straighter tracks, at ages 4-dpf (estimate = - 0.970 ± 0.210, z = -4.614, *p* < 0.001), 5-dpf (estimate = -0.866 ± 0.211, z = -4.102, *p* < 0.001), and 6-dpf (estimate = -0.852 ± 0.211, z = -4.029, *p* < 0.001). Pairwise comparisons revealed that tortuosity of larval tracks was significantly higher on the F/2 control slips compared to *G. marina* slips at 4-dpf (Fig. 3, estimate = 0.391 ± 0.188, z = 2.077, *p* < 0.05), 5-dpf (estimate = 0.535 ± 0.187, z = 2.856, *p* < 0.01), and 6-dpf (estimate = 0.517 ± 0.188, z = 2.746, *p* < 0.01). Track tortuosity was also significantly lower on F/2 control slips from 2-dpf until 4-dpf, at 5- and 6-dpf, tortuosity was no different between areas. Track tortuosity was also significantly lower on the *G. marina* coated slips compared to the no-slip side of the *G. marina* chamber from 3-dpf onwards.

**Figure 3.**
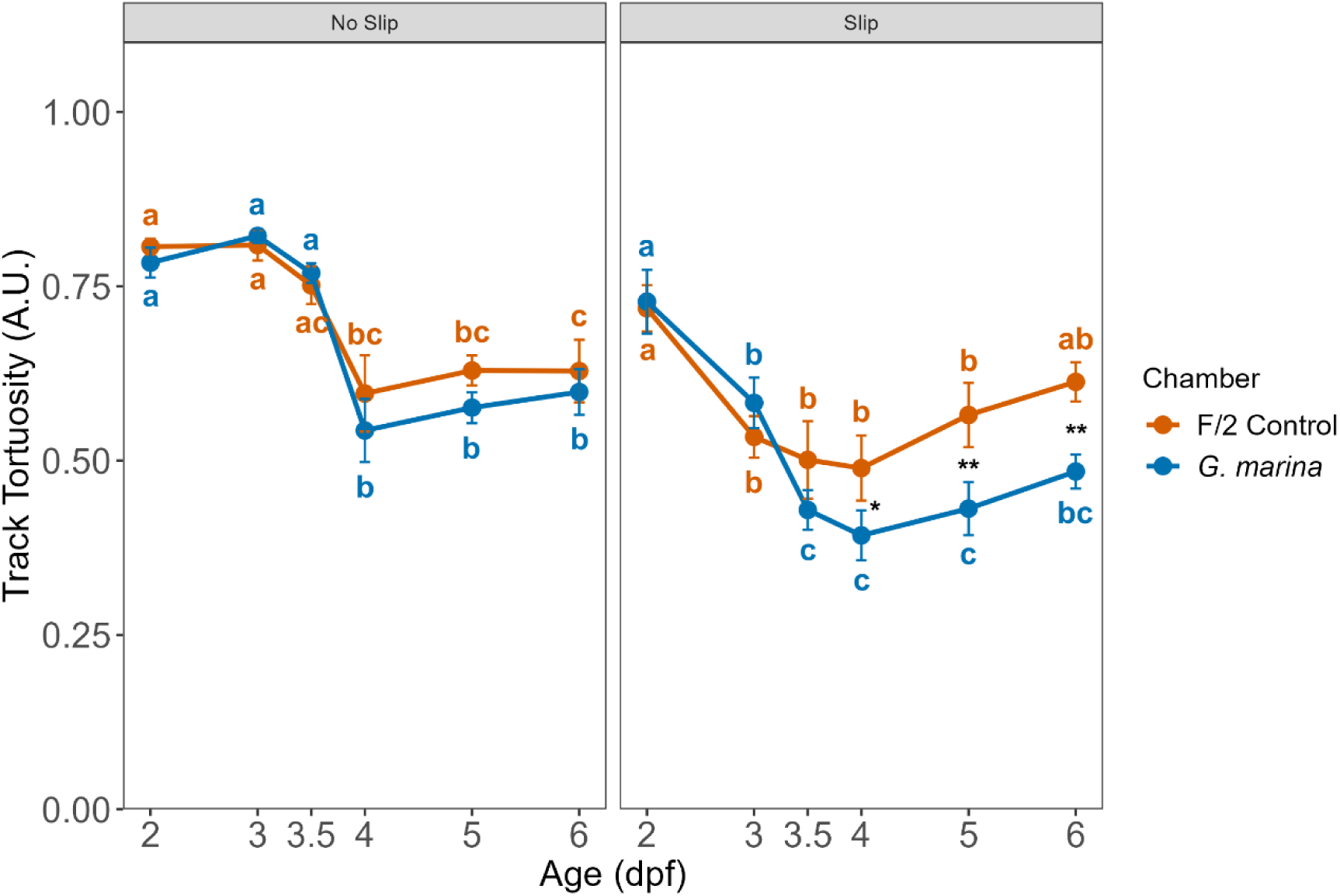
The effect of the *G. marina* biofilm on track tortuosity by age and area. Track tortuosity of all larval tracks on both the no-slip area (left panel) and slip area (right panel) of the chambers are shown for the F/2 control chamber (orange) and *G. marina* chamber (blue) across ages. The ages of *P. dumerilii* larvae are expressed as days post-fertilisation (dpf). Different letters of the same colour represent statistically significant differences within chambers and areas across ages. Asterisks indicate statistically significant differences between treatments within ages and areas where * *p* < 0.05, ** *p* < 0.01. *N* = 6 videos for each age group.

As the coverage of the *G. marina* biofilm on the slips was variable, the effect of diatom coverage on the proportion of crawling larvae, crawling speed, and track tortuosity was explored. Here, there were no significant correlations of these behaviours with *G. marina* cell coverage at any age (Supplementary Figure 2). Therefore, diatom coverage did not have a significant impact on settlement behaviour within the range of biofilm densities that were tested.

### Larval settlement

To assess the rate at which *P. dumerilii* larvae settle on the F/2 control and *G. marina* coated slips over time, linear models were fitted to the RSS data for each age group. Larvae aged between 3.5- and 5-dpf did not show any kind of settlement response to the control slip, with their numbers either remaining the same throughout the experiment or decreasing over time (Fig. 4A). At 3-dpf, the numbers of *P. dumerilii* larvae associated with the control slip increased slowly over the course of the experiment. The numbers for 2-dpf and 6-dpf larvae also increased slowly, but not to the same extent as on the *G. marina* biofilm.

**Figure 4.**
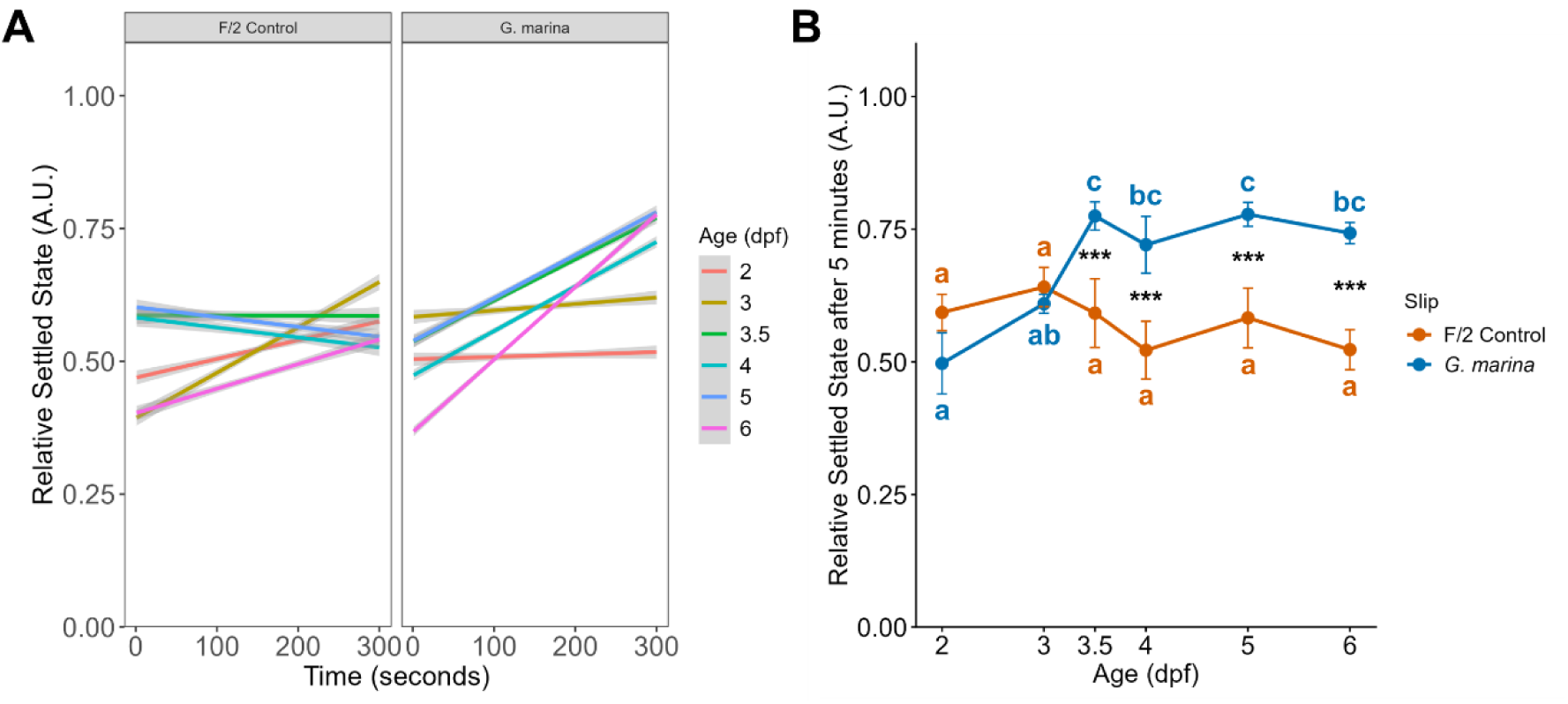
The Relative Settled State (RSS) of *P. dumerilii* larvae on the F/2 control slips or *G. marina* slips. (A) The RSS of larvae at different ages over the 5-minute period of each experiment with lines plotted using a multiple linear regression model. (B) The RSS of each age group on F/2 control slips and *G. marina* slips at the end of the 5-minute recordings. In (B) different letters of the same colour represent statistically significant differences within treatments across ages. Asterisks indicate statistically significant differences between treatments within ages where *** *p* < 0.001. *N* = 6 videos for each age group.

For the *G. marina* coated slips 2- and 3-dpf larvae were observed to interact with the slip within the first seconds of the recording, with the interaction occurring faster compared to the control slip (Fig. 4A). However, their relative numbers remain constant indicating that larvae did not remain closely associated with the slip for long or simply swam over the biofilm before leaving the ROI. It is possible that 3-dpf larvae could crawl over the biofilm, but they appear to swim away after a short period of contact. There was an increase in the relative number of larvae on the *G. marina* slip at ages 3.5-, 4-, 5, and 6-dpf (Fig. 4A). Increases in RSS over time indicates that larvae at these ages remained on the biofilm once contact had been made. Traces of the raw data plotted over the course of the recordings show short, momentary increases in RSS due to larvae swimming into the ROI for a few seconds before leaving (Supplementary Figure 3). Larvae at ages 3.5-to 6-dpf also colonise the *G. marina* biofilm in the highest numbers.

To quantify if these rates of change in RSS were statistically significant, the slopes of the linear models were compared. The steepness of the slopes was used as a proxy for the rate of RSS change over time, with steeper slopes indicating more rapid colonisation of the biofilm. The mean RSS slope steepness for 3.5-, 4-, 5-, and 6-dpf larvae on the *G. marina* biofilm was 0.000777, 0.000839, 0.000810, and 0.00136, respectively (Fig. 4A). There was an effect of larval age on the RSS slopes (F_5,30_ = 3.606, *p* < 0.05). However, pairwise comparisons of the slopes revealed that there were no significant differences in the steepness of the slopes between 3.5-, 4-, 5-, and 6-dpf larvae. Only the difference in the slopes between 2-dpf and 6-dpf, and 3-dpf and 6-dpf were statistically significant (Supplementary Figure 4). Therefore, despite initially starting from a lower number on the biofilm, the rate of change in RSS for 6-dpf larvae seems to be quicker than 3.5-, 4-, and 5-dpf larvae, but not significantly so.

The interactive effects of age and slip type on RSS at the end of the 5 min recordings were assessed by GLM. The model revealed that there were significant interactions between the *G. marina* coated slip on RSS at ages 3.5-dpf (estimate = 1.224 ± 0.340, z = 3.600, *p* < 0.001), 4-dpf (estimate = 1.266 ± 0.334, z = 3.792, *p* < 0.001), 5-dpf (estimate = 1.257 ± 0.340, z = 3.692, *p* < 0.001), and 6-dpf (estimate = 1.318 ± 0.335, z = 3.935, *p* < 0.001), compared to the RSS of 2-dpf larvae on F/2 control slips (Supplementary Table 4). Pairwise comparisons also revealed that the RSS of larvae was significantly lower on F/2 control slips compared to *G. marina* coated slips at 3.5-dpf (estimate = -0.839 ± 0.250, z = -3.354, *p* < 0.001), 4-dpf (estimate = -0.880 ± 0.241, z = -3.646, *p* < 0.001), 5-dpf (estimate = -0.871 ± 0.250, z = -3.479, *p* < 0.001), and 6-dpf (estimate = -0.932 ± 0.243, z = -3.837, *p* < 0.001; Fig 4B). There were no differences in RSS between ages within the F/2 control treatment. Within the *G. marina* treatment, RSS significantly increased between 3- and 3.5-dpf which remained elevated with no significant differences in RSS detected from 3.5-dpf to 6-dpf (Fig. 4B).

### Effect of *G. marina* biofilm on swimming and crawling trajectories

We questioned if the increasing RSS seen on the *G. marina* slip may be due to older larvae changing the way they swim or crawl. We hypothesised that if the biofilm released volatile cues, this would reduce swimming tortuosity (make swimming straighter), and increase swimming speeds, reducing the time taken to find the biofilm. To do this, we subset swimming and crawling tracks of all larvae in each chamber into separate datasets and calculated the tortuosity of these swimming and crawling tracks. Our analysis shows that there was no interactive effect between age and slip type on swimming or crawling tortuosity. However, for swimming tracks, there was a significant effect of age on their straightness (Fig. 5A, F_5,64_ = 28.92, *p* < 0.001) and that swimming tortuosity significantly reduced at 4-dpf (*p* < 0.001), 5-dpf (p < 0.001), and 6-dpf (*p* < 0.001). There was no effect of F/2 control or *G. marina* biofilm on swimming tortuosity (Fig. 5B). There was also a significant effect of age on the speed of swimming tracks (Fig. 5C, F_1,139_ = 23.757, *p* < 0.001) with swimming speeds increasing from 3-dpf onwards, but there was no effect of treatment (Fig. 5D). For crawling tracks, there was also no interactive effect of age and treatment, nor was there an effect of age on their tortuosity (Fig. 5E). The presence of *G. marina* biofilm did reduce crawling tortuosity across all ages compared to the controls (Fig. 5F, t_56_ = 2.014, *p* < 0.05). The speed of crawling tracks did increase with age (Fig. 5G, F_1,116_ = 19.01, *p* < 0.001) which became significant at 4-dpf. The presence of the *G. marina* biofilm significantly decreased crawling speeds across all ages (Fig. 5H, t_113_ = 3.767, *p* < 0.001).

**Figure 5.**
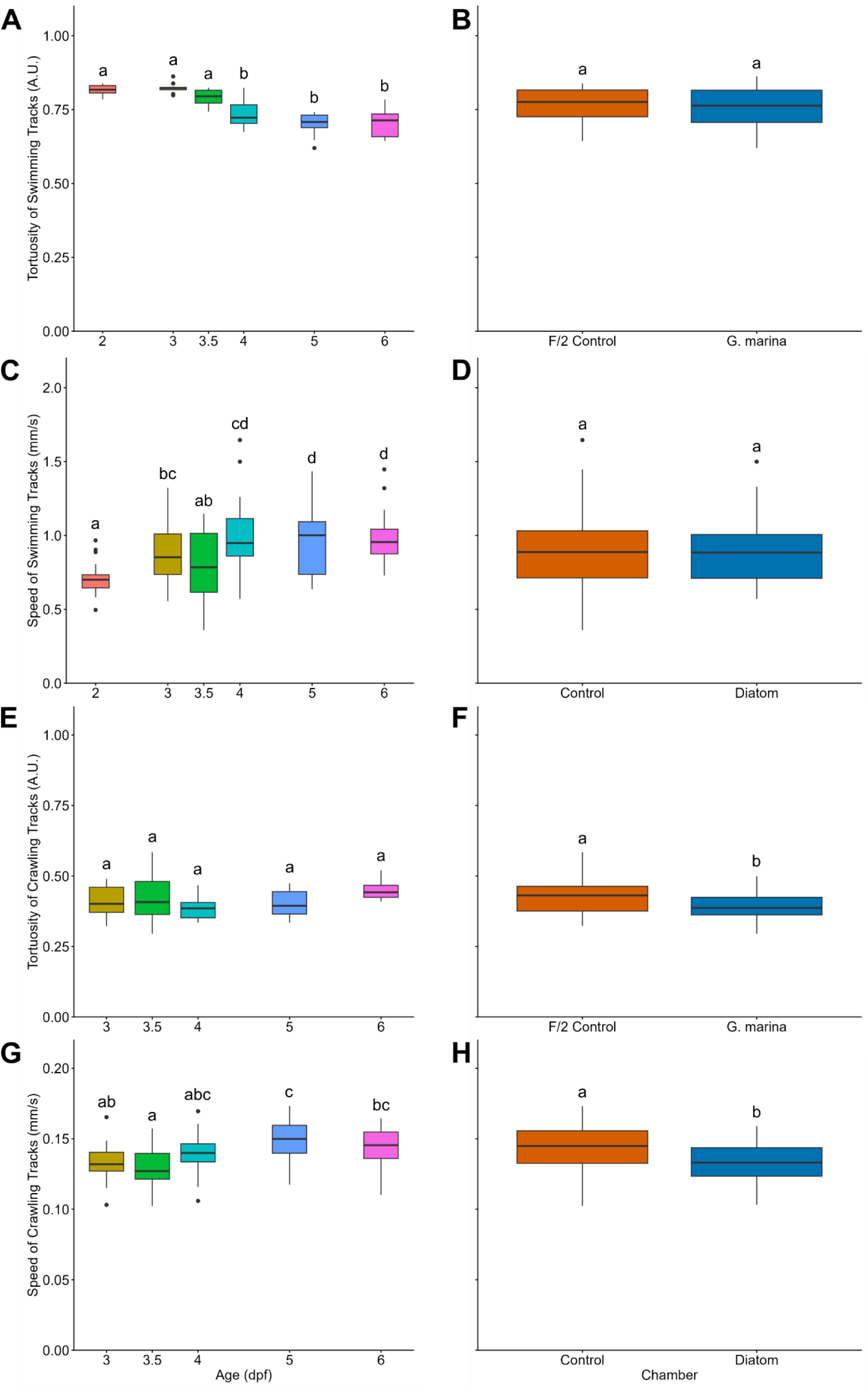
Tortuosity and speed of swimming and crawling tracks. (A & B) The effect of age and treatment on swimming tortuosity. (C & D) The effect of age and treatment on swimming speeds. (E & F) The effect of age and treatment on crawling tortuosity. (G & H) Effect of age and treatment on crawling speeds. Data for 2-dpf larvae was removed from E-H as larvae at this age do not crawl. Each box represents the interquartile range with the median represented by a solid horizontal line. The whiskers extending from each box represents the upper and lower ranges with black points indicating outliers. Letters above boxes indicate statistically significant differences. *N* = 6 videos for each age group.

### Settlement preference choice

*P. dumerilii* larvae were also given a choice to settle on an F/2 control slip or a *G. marina* coated slip by presenting the two surfaces within the same environment, rather than being separated within their own chamber. This revealed a highly significant interaction between the effects of age and slip type on larval settlement behaviour (F_5,58_ = 8.098, *p* < 0.001). More larvae settled on the diatom biofilm compared to the F/2 control in the same chamber at ages over 3.5-dpf (Fig. 6). Pairwise comparisons of settlement behaviour show that there are highly significant differences in the proportion of larvae settled on the *G. marina* slip compared to the F/2 control slip at ages 4- (*p* < 0.001), 5- (*p* < 0.001), and 6-dpf (*p* < 0.001). Preference behaviour starts to become evident at 3.5-dpf, but this difference is not statistically significant. There were also no differences in the proportion of settled larvae at 2- and 3-dpf, with almost identical proportions between F/2 control and diatom biofilm slips. At least half of the larvae in the cuvette did not settle on a slip during the 5 min filming time. For 4-, 5-, and 6-dpf old worms the proportion of larvae in the cuvette not on a slip equates to approximately 50%, 55%, and 56% respectively.

**Figure 6.**
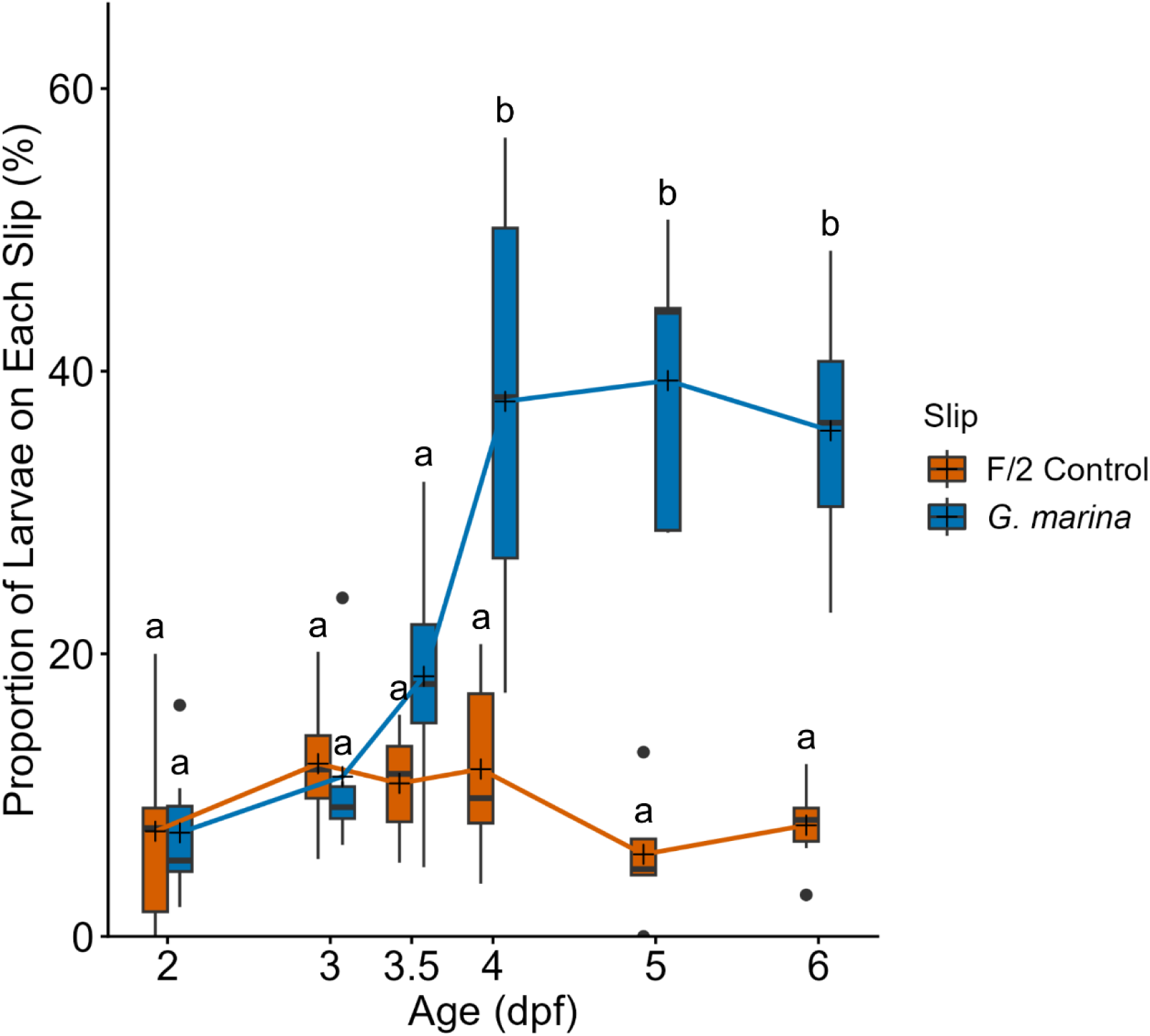
Settlement preference behaviour of *P. dumerilii* larvae towards F/2 control slips or *G. marina* slips expressed as a proportion (%) of the total larvae within the cuvette. Ages of larvae are expressed in days post-fertilisation (dpf). Each box represents the interquartile range with the median represented by a solid horizontal line and the mean represented by crosses. Whiskers represent the upper and lower ranges with black points indicating outliers. Different letters above the bars indicate statistically significant differences. *N* = 6 for each age group.

## Discussion

In this study, we investigated how settlement behaviour of *P. dumerilii* larvae changes over the course of development in the presence and absence of a diatom biofilm of *Grammatophora marina*. By developing a larval tracking and analysis pipeline several key behavioural changes that occur during biofilm exploration were established, which include increases in crawling, reduced crawling speeds, straighter crawling, increased response to settlement cues and increased association with the biofilm, the level of which changes depending on larval age. Younger larvae tend not to show any settlement preference behaviour, whereas larvae over 3.5-dpf demonstrate increasingly altered behaviour as a result of biofilm interaction. From these analyses, we can better understand the behavioural repertoire of developing *P. dumerilii,* and begin to identify settlement behaviours of this species in different environmental contexts. Moreover, this study validates a methodology for accurately characterising settlement behaviour in marine invertebrate larvae, that is also transferable to a wide range of other experimental models.

### Ontogeny of *P. dumerilii* settlement behaviour

Our results show a graded change from swimming to crawling behaviour in response to microalgal biofilms throughout *P. dumerilii* development. Larvae must first swim to and then contact the biofilm, where crawling behaviour becomes more common than swimming. Crawling speeds on the biofilm are slower than when biofilm is absent, but these increase with age. Tracks become straighter when crawling on the biofilm compared to when swimming over it. Finally, older larvae are more likely to stay on the biofilm (Fig. 7). Having established these behavioural traits, these specific behaviours can be incorporated in future studies focussing on how other biotic factors, such as dissolved chemical cues, influence settlement behaviour. As these experiments were performed in still water, it would be useful in the future to test how flow rates and turbulent flow may affect *P. dumerilii* behavioural responses to *G. marina* biofilms. A recent study on the effect of flow on biofilm exploration behaviour of *H. elegans* larvae showed that although flow can alter swimming trajectories, it did not alter the percentage of larvae touching a biofilm (Koehl et al., 2022), suggesting that the predominant behavioural responses seen in our study will also be maintained under flow.

**Figure 7.**
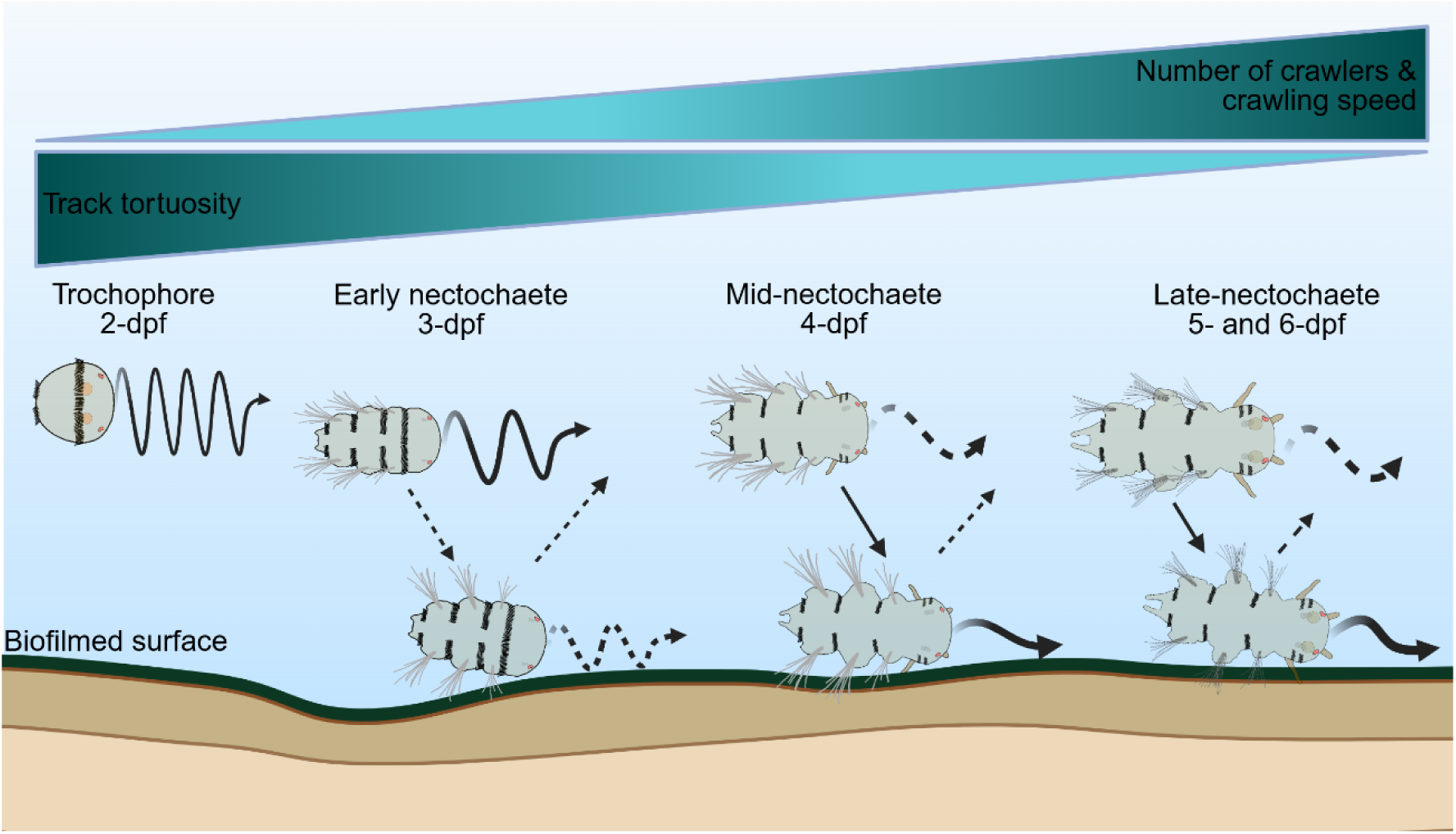
Behavioural responses of *Platynereis dumerilii* larvae to *G. marina* biofilms throughout development. Two-day old trochophore larvae are mostly unresponsive to the biofilm, swimming over it with a helical swimming pattern. At 3-days old, larvae may begin surface exploration and crawling starts, but the larvae are more likely to continue swimming or swim away after biofilm contact is made. From 3.5- to 4-days old onwards, sinking out of the water column is much more common. Larvae at these ages will initiate crawling once contact with the biofilm is made, their crawling becomes straighter compared to when they were swimming over the biofilm, and larvae are increasingly likely to remain on the biofilm rather than swimming away. Solid lines represent the most likely mode of movement, swimming or crawling, at those ages when a biofilm is present. Dashed lines represent modes of movement that are less likely at those ages when a biofilm is present. The thickness of the lines represents movement speed, with thicker lines representing faster movement. Cartoon representations of *Platynereis* larvae were adapted from Fischer et al., (2010). Created with BioRender.com.

### Biofilm surface exploration in *P. dumerilii* larvae

*P. dumerilii* larvae exhibit surface exploration behaviour from 3.5- to 4-dpf, indicating settlement competency begins around this time in development. Once larvae at these ages encounter a *G. marina* biofilm, they switch from swimming to mostly crawling, reduce their movement speed, their tracks become straighter compared to when they are swimming, and the larvae stay in contact with the biofilm for longer compared to controls. Similar behavioural sequences of settlement on microbial biofilms that include reduced movement speeds followed by touching and attachment have been recorded in competent larvae of the polychaete *Hydroides elegans* (Hadfield et al., 2021) and the barnacle *Amphibalanus* (formally *Balanus*) *amphitrite* (Maleschlijski et al., 2015; Miron et al., 2000). There were consistently greater numbers of larvae moving with tracks with a greater number of turns on the no-slip side of the dual-chamber cuvette when compared to the side with the biofilm or control slips. It has been suggested that larval locomotion has curved or helical tracks when searching for a settlement cue, or when there is no settlement cue to be detected (Hadfield et al., 2014; Maciejewski et al., 2019; Maleschlijski et al., 2015). Whether the higher tortuosity that was seen on the no-slip side of the chamber is a searching behaviour in *P. dumerilii* is uncertain and it is likely just the inherent swimming pattern of the larvae. *P. dumerilii* trochophore larvae have a spiralling swimming pattern (Jékely et al., 2008) which would score highly for tortuosity. Although older larvae swim with a torpedo-like body shape, they will use their trunk muscles to create left and right turns as they swim (Randel et al., 2014). However, a decrease in tortuosity on the *G. marina* biofilm, and therefore a slight increase in the track straightness, could be a way for *P. dumerilii* to maximise the total area of the biofilm they can explore before making the decision to settle.

### Inducive elements of *G. marina* biofilm

Once surface exploration has been initiated, or when the larvae have determined that the biofilm represents a suitable settlement cue, *P. dumerilii* at 3.5-dpf onwards will remain in contact with the biofilm. This is shown by the positive relationship with RSS over time, as well as the preferential settlement of *P. dumerilii* on the *G. marina* biofilm when also presented with a no biofilm control. Once contact with the biofilm is made, there are likely both chemical and physical signals presented to the larvae that they can sense and respond to. It is well established that microbial biofilms induce settlement in marine larvae, often in a species-specific manner. A common feature of marine diatom biofilms is the production of extracellular polymeric substances (EPS), rich in polysaccharides, which form a major component of the biofilm (Hoagland et al., 1993; Laroche, 2022; Li et al., 2023; Silva et al., 2024). EPS is a likely to be one possible cue that *P. dumerilii* is using to determine a suitable settlement location. The EPS extracted from diatom biofilms have been shown to increase settlement in *A. amphitrite* (Patil & Anil, 2005), *H. elegans* (Lam et al., 2005), and the clam *Macoma balthica* (van Colen et al., 2009). There remain some questions surrounding whether larval contact with biofilms is necessary for initiating settlement, or if biofilms release soluble chemo-attractants. Larvae of the annelid,*H. elegans*, neither settle on unfilmed surfaces set opposite to a biofilm (Cindy et al., 2003), nor when separated above from a biofilm by a mesh screen (Hadfield et al., 2014). Settlement rates have been shown to be lower for larvae of *A. amphitrite* when exposed to EPS extracted from seawater in which diatom biofilms had been maintained than from EPS extracted directly from the biofilm (Patil & Anil, 2005). A necessity for physical contact with the inductive cue has also been demonstrated for settlement in *A. amphitrite,* where treating biofilms with lectins blocked surface-bound sugars from being available to larvae, reduced settlement rates (Jouuchi et al., 2007). Thus, whilst no testing was undertaken to detect chemical signatures in the seawater in our study, based on previous research and a lack of behavioural changes demonstrated in larvae prior to contacting the biofilms in our study, we propose that settlement cues are most likely to be predominantly available to *P. dumerilii* on the surface of the *G. marina* biofilm.

Although the relative number of *P. dumerilii* larvae increased on the biofilms over time, it is not clear that this was due to chemotaxis, and larvae did still locate and interact with control slips for a short time when they are unlikely to be producing any volatile settlement cues. We also saw no effect of the *G. marina* biofilm on the tortuosity and speed of swimming tracks which suggests that there were limited dissolved cues in the water column to which the larvae were actively responding. Nonetheless, with the current experiments, we cannot rule out the possibility that both chemotaxis and physical contact both contribute to changes in *P. dumerilii* settlement behaviour from 3.5-dpf onwards.

It has been well documented that some marine larvae are able to respond to dissolved chemical cues resulting in settlement behaviours (Da-Anoy et al., 2017; Hadfield & Koehl, 2004; Jensen & Morse, 1990; Meyer-Kaiser et al., 2019; Pawlik & Faulkner, 1986; Turner et al., 1994). Thus, to confirm whether *P. dumerilii* must physically encounter the biofilm to induce settlement behaviours, and rule out the possibility of soluble chemicals contributing to this process, further study is required.

### Hierarchy of sensory modalities during *Platynereis dumerilii* development

As larvae develop and their sensory organs and nervous systems become more mature, the importance of different sensory inputs and their effect on larval behaviour also changes (Hodin et al., 2017).. For an early larva, it may be more important to gauge light and pressure to determine their vertical position in the water column. As the larva ages and reaches competency, the larva may then prioritise the detection of food or conspecifics to locate suitable settlement locations. For *P. dumerilii*, trochophore to early nectochaete stages (1-3 days old) possibly prioritise light detection, phototaxis, and barotaxis to select an appropriate vertical position in the ocean (Bezares Calderón et al., 2024; Gühmann et al., 2015; Jékely et al., 2008; Verasztó et al., 2018). From 3.5-days old, when settlement appears to begin, the larva may incorporate more mechanoreception, aiding exploration of the seafloor (Bezares-Calderón et al., 2018). Finally, from mid- to late-nectochaete (4-6 days old), chemoreception may dominate as the larva looks to locate suitable settlement cues within biofilms that represent a food source and substrate for settlement (Chartier et al., 2018; Hird et al., 2024). To confirm this hypothesis, further investigation of the chemo-sensation in early larval stages (trochophore to early nectochaete), in terms of both the suite of chemical cues detected and responsive behaviours, is required. The scale over which each of these sensory inputs become available to the larvae will also influence their priority for the larvae. These experiments were conducted at a scale of millimetres to centimetres which is the hypothesised scale at which chemical cues would be dominant (Hodin et al., 2017).

## Conclusion

Larval *P. dumerilii* exhibit developmental stage-specific behaviours in response to their environment, but also a developmentally programmed inherent transition from swimming to crawling, with higher proportions of larvae crawling with age even in the absence of biofilm. In the presence of *G. marina* biofilms, 3.5-dpf and older *P. dumerilii* larvae will switch from swimming to crawling when responding to the biofilm or when physical contact is made. When exploring the biofilm, crawling larvae travel in straighter trajectories. The proportions of larvae performing such behaviours and the magnitude of their behavioural responses change with age, likely reflecting enhanced sensory capabilities and changes in sensory cue hierarchies through larval development. Our study has generated a baseline of short-term behavioural responses of different ages of larvae to diatom biofilms. From 3.5-dpf onwards, *P. dumerilii* are also able to distinguish between similar physical surfaces with and without preferred settlement cues when presented with both options. The ability of the animals to make this distinction early in development will have positive impacts on the fitness of the animal later in life.

The analytical pipeline developed in this study, along with our characterisation of the baseline behavioural responses of wildtype *P. dumerilii* larvae to biofilm at different ages, enables future studies of the influence of other abiotic parameters including flow, temperature, and salinity on larval behaviour in this species. These tools also provide the groundwork for further investigations of the role of the nervous system in larval-biofilm interactions through characterisation of the behavioural response of mutant larvae lacking different neurotransmitters to *G. marina* biofilms. Finally, the simplicity of the recording design is easily transferable to the tracking of larvae of other species of annelids, molluscs, and crustaceans, for comparative studies of the evolutionary relationship between marine larvae and benthic biofilms and characterisation of the diversity of marine larval settlement behaviour.

## Acknowledgments

We thank Adam Johnstone and Richard Silcox for their contributions in running the *Platynereis dumerilii* culture at the Marine Invertebrate Culture Unit (MICU) at the University of Exeter. We also thank Luis Bezares-Calderon and Emelie Brodrick for helpful directions and discussions on larval behaviour recording and analysis in R Studio.

## Author Contributions

C.T.: conceptualisation, data curation, formal analysis, investigation, methodology, project administration, writing – original draft, writing – review and editing; S.V.: writing – review and editing, methodology; R.P.E.: writing – review and editing, methodology, formal analysis; E.A.W.: conceptualisation, methodology, formal analysis, funding acquisition, project administration, writing – review and editing.

