## Supplementary File 1 for "Ontogeny of settlement behaviours in response to *Grammatophora marina* diatom biofilms in the marine polychaete, *Platynereis dumerilii*"

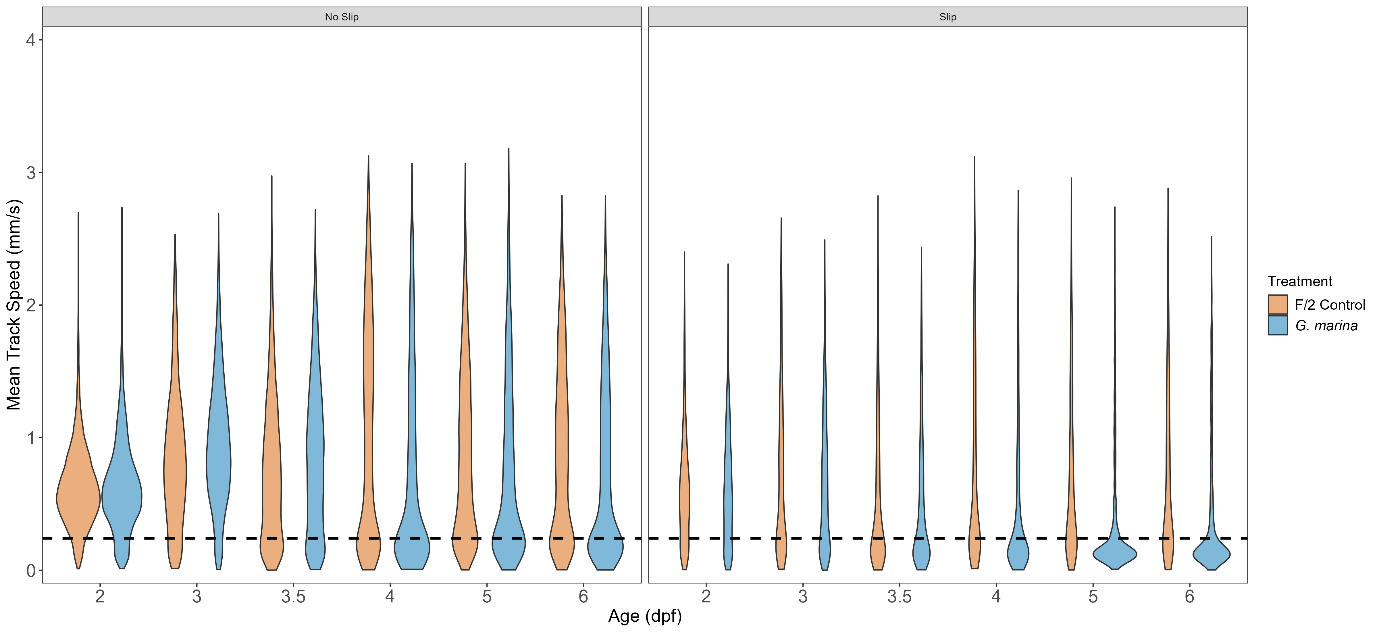

Supplementary Figure 1. Mean larval track speeds. Violins show the distribution of the mean speed data between chambers containing an F/2 control slip (orange) and *G. marina* biofilm slip (blue). The data is split between tracks from the “No Slip” (left panel) and “Slip” (right panel) areas of each chamber. The dashed line is set to 0.24 mm/s representing the cutoff for the crawling speed data.

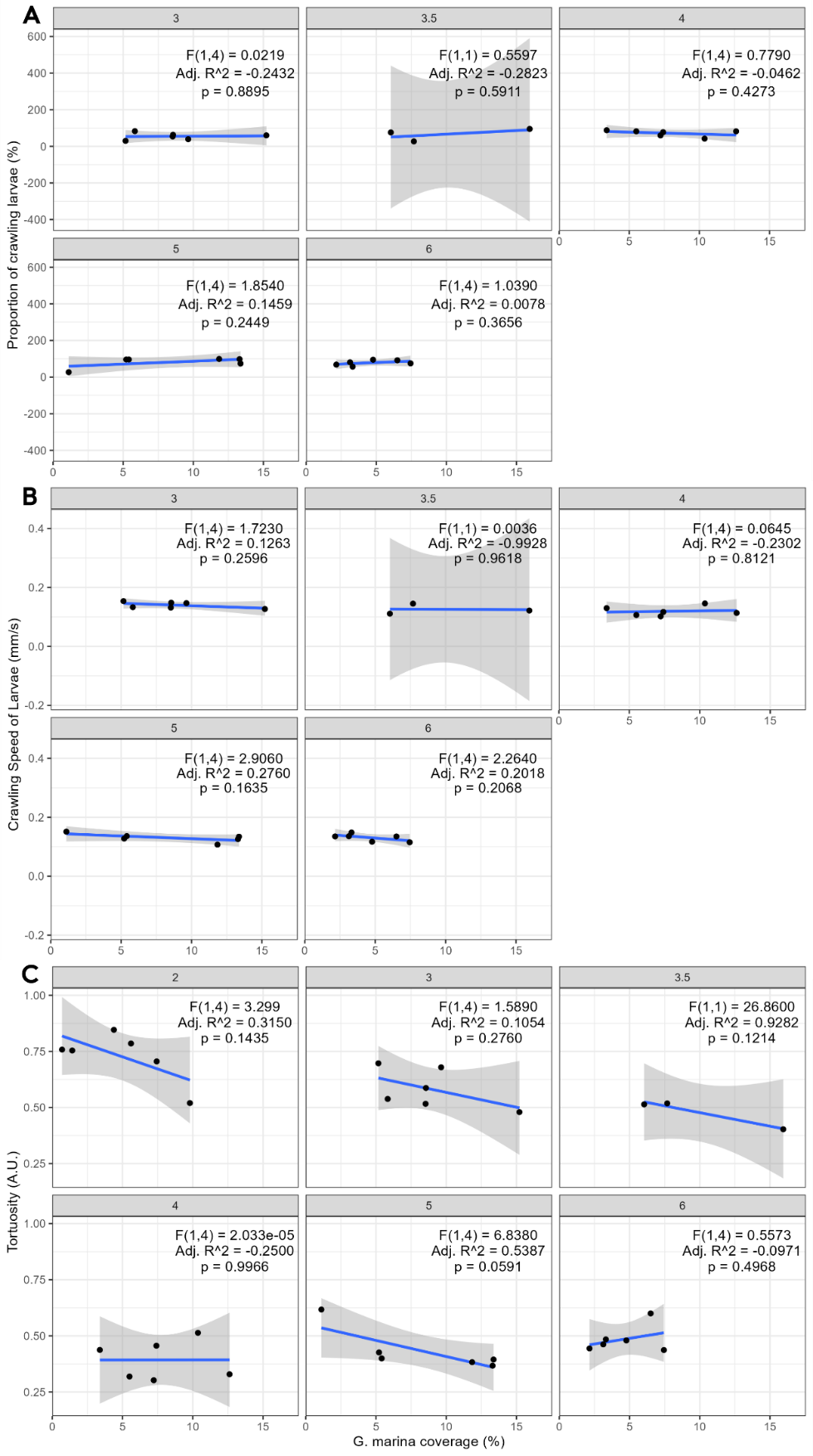

Supplementary Figure 2. Relationships between the coverage of the *G. marina* biofilm on the slips with (A) the proportion of crawling larvae, (B) the crawling speed of larvae, and (C) track tortuosity. The age in days post-fertilisation (dpf) is shown in the title of each plot. Data for 2-dpf data has been removed from (A) as larvae at this age do not crawl. Each blue line represents the fit of the linear models and grey ribbons represent ± standard deviation. *N* = 6 slips except for data relating to 3.5-dpf where *n* = 3 slips.

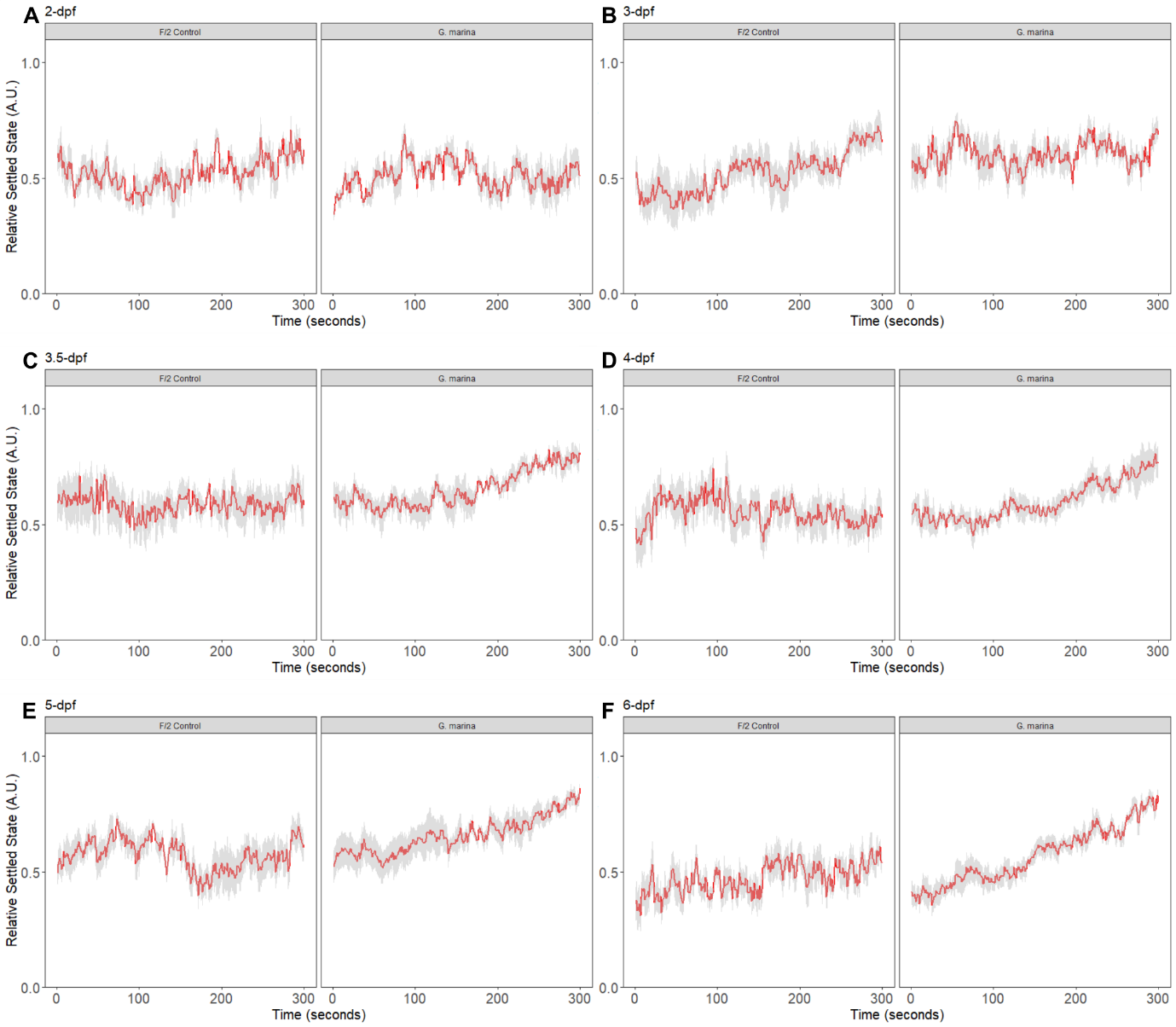

Supplementary Figure 3. Raw RSS data plotted per second over five minutes. For A-F, the left panel shows RSS of larvae on the F/2 control slip, and the right panel shows the RSS on the *G. marina* slip. The red trend line represents the mean RSS per second, and the grey ribbon represents ± standard error. *N* = 6 per age group.

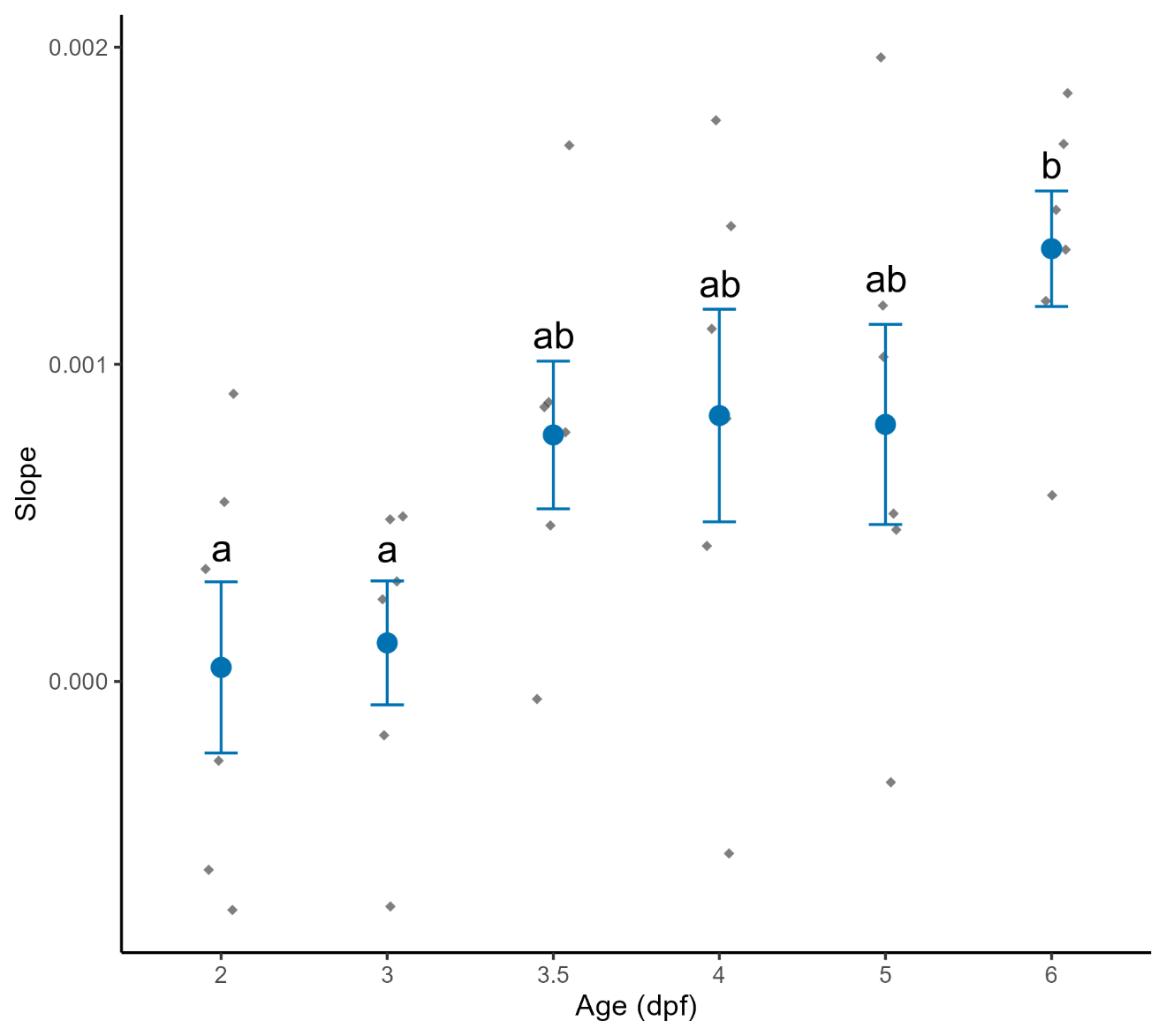

Supplementary Figure 4. Slopes retrieved from the linear models fitted to the RSS data for the *G. marina* coated slips per age group. Data is shown as the mean ± standard error with raw data points overlaid as grey diamonds. Different letters above the data points indicates significant differences. *N* = 6 per age group.

Supplementary Table 1. Output table from the beta GLM analysis of the effects of Age, Treatment, and Area on the proportion of crawling larvae in the two-chamber assay. Asterisks indicate significant effects compared to the intercept (Age 3, F/2 control, no slip area), * *p* < 0.05, ** *p* < 0.01, *** *p* < 0.001.

| Model | Fixed effects | Estimate | Standard error | z-value | *p*-value |  |
| --- | --- | --- | --- | --- | --- | --- |
| Proportion of crawling larvae | | | | | | |
| Age * Treatment + Area | (Intercept) | -1.606 | 0.211 | -7.611 | 2.72e-14 | *** |
|  | Age 3.5 | 0.557 | 0.273 | 2.042 | 0.041 | * |
|  | Age 4 | 0.285 | 0.280 | 1.016 | 0.310 |  |
|  | Age 5 | -0.024 | 0.285 | -0.085 | 0.932 |  |
|  | Age 6 | 0.257 | 0.281 | 0.914 | 0.361 |  |
|  | Treatment (Diatom) | -0.357 | 0.289 | -1.234 | 0.217 |  |
|  | Area (Slip) | 0.912 | 0.124 | 7.385 | 1.52e-13 | *** |
|  | Age 3.5 * Diatom | 0.199 | 0.392 | 0.507 | 0.612 |  |
|  | Age 4 * Diatom | 1.258 | 0.396 | 3.181 | 0.001 | ** |
|  | Age 5 * Diatom | 1.423 | 0.397 | 3.582 | 0.0003 | *** |
|  | Age 6 * Diatom | 1.215 | 0.396 | 3.070 | 0.002 | ** |

Supplementary Table 2. Output table from the gaussian GLM analysis of the effects of Age, Treatment, and Area on larval crawling speed in the two-chamber assay. Asterisks indicate significant effects compared to the intercept (Age 3, F/2 control, no slip area), * *p* < 0.05, *** *p* < 0.001.

| Model | Fixed effects | Estimate | Standard error | t-value | *p*-value |  |
| --- | --- | --- | --- | --- | --- | --- |
| Larval crawling speeds | | | | | | |
| Age * Treatment + Area | (Intercept) | 0.139 | 0.004 | 35.288 | <2e-16 | *** |
|  | Age 3.5 | -0.006 | 0.005 | -1.090 | 0.278 |  |
|  | Age 4 | 0.011 | 0.005 | 2.055 | 0.042 | * |
|  | Age 5 | 0.021 | 0.005 | 4.016 | 0.0001 | *** |
|  | Age 6 | 0.014 | 0.005 | 2.702 | 0.008 |  |
|  | Treatment (Diatom) | -0.005 | 0.005 | -0.948 | 0.345 |  |
|  | Area (Slip) | -0.008 | 0.002 | -3.432 | 0.0008 | *** |
|  | Age 3.5 * Diatom | 0.002 | 0.008 | 0.299 | 0.765 |  |
|  | Age 4 * Diatom | -0.010 | 0.008 | -1.376 | 0.171 |  |
|  | Age 5 * Diatom | -0.011 | 0.008 | -1.475 | 0.143 |  |
|  | Age 6 * Diatom | -0.008 | 0.008 | -1.101 | 0.273 |  |

Supplementary Table 3. Output table from the beta GLM analysis of the effects of Age, Treatment, and Area on track tortuosity in the two-chamber assay. Asterisks indicate significant effects compared to the intercept (Age 2, F/2 control, no slip area), * *p* < 0.05, ** *p* < 0.01, *** *p* < 0.001.

| Model | Fixed effects | Estimate | Standard error | z-value | *p*-value |  |
| --- | --- | --- | --- | --- | --- | --- |
| Tortuosity of larval tracks | | | | | | |
| Age * Treatment * Area | (Intercept) | 1.386 | 0.162 | 8.561 | <2e-16 | *** |
|  | Age 3 | 0.032 | 0.230 | 0.142 | 0.888 |  |
|  | Age 3.5 | -0.285 | 0.221 | -1.292 | 0.196 |  |
|  | Age 4 | -0.970 | 0.210 | -4.614 | 3.96e-6 | *** |
|  | Age 5 | -0.866 | 0.211 | -4.102 | 4.10e-5 | *** |
|  | Age 6 | -0.852 | 0.211 | -4.029 | 5.59e-5 | *** |
|  | Treatment (Diatom) | -0.117 | 0.225 | -0.519 | 0.604 |  |
|  | Area (Slip) | -0.452 | 0.217 | -2.080 | 0.036 | * |
|  | Age 3 * Diatom | 0.180 | 0.324 | 0.554 | 0.579 |  |
|  | Age 3.5 * Diatom | 0.188 | 0.311 | 0.604 | 0.546 |  |
|  | Age 4 * Diatom | -0.121 | 0.294 | -0.413 | 0.680 |  |
|  | Age 5 * Diatom | -0.102 | 0.295 | -0.348 | 0.728 |  |
|  | Age 6 * Diatom | -0.020 | 0.302 | -0.069 | 0.945 |  |
|  | Age 3 * Slip | -0.830 | 0.295 | -2.748 | 0.006 | ** |
|  | Age 3.5 * Slip | -0.632 | 0.287 | -2.140 | 0.032 | * |
|  | Age 4 * Slip | -0.009 | 0.287 | -0.031 | 0.976 |  |
|  | Age 5 * Slip | 0.195 | 0.288 | 0.677 | 0.498 |  |
|  | Age 6 * Slip | 0.374 | 0.290 | 1.292 | 0.196 |  |
|  | Diatom * Slip | 0.186 | 0.306 | 0.609 | 0.542 |  |
|  | Age 3 * Diatom * Slip | -0.050 | 0.428 | -0.117 | 0.907 |  |
|  | Age 3.5 * Diatom * Slip | -0.555 | 0.418 | -1.329 | 0.184 |  |
|  | Age 4 * Diatom * Slip | -0.339 | 0.405 | -0.836 | 0.403 |  |
|  | Age 5 * Diatom * Slip | -0.502 | 0.406 | -1.238 | 0.216 |  |
|  | Age 6 * Diatom * Slip | -0.566 | 0.407 | -1.392 | 0.164 |  |

Supplementary Table 4. Output table from the beta GLM analysis of the effects of Age and Treatment on the Relative Settled State (RSS) of larvae in the two-chamber assay. Asterisks indicate significant effects compared to the intercept (Age 2, F/2 control, no slip area), * *p* < 0.05, *** *p* < 0.001.

| Model | Fixed effects | Estimate | Standard Error | z-value | *p*-value |  |
| --- | --- | --- | --- | --- | --- | --- |
| Relative Settled State (RSS) | | | | | | |
| Age * Treatment | (Intercept) | 0.371 | 0.164 | 2.261 | 0.024 | * |
|  | Age 3 | 0.196 | 0.235 | 0.834 | 0.405 |  |
|  | Age 3.5 | -0.009 | 0.232 | -0.038 | 0.970 |  |
|  | Age 4 | -0.284 | 0.231 | -1.230 | 0.219 |  |
|  | Age 5 | -0.024 | 0.232 | -0.105 | 0.916 |  |
|  | Age 6 | -0.281 | 0.231 | -1.128 | 0.223 |  |
|  | Slip (Diatom) | -0.386 | 0.230 | -1.673 | 0.094 |  |
|  | Age 3 * Diatom | 0.248 | 0.329 | 0.754 | 0.451 |  |
|  | Age 3.5 * Diatom | 1.224 | 0.340 | 3.600 | 0.0003 | *** |
|  | Age 4 * Diatom | 1.266 | 0.334 | 3.792 | 0.0002 | *** |
|  | Age 5 * Diatom | 1.257 | 0.340 | 3.692 | 0.0002 | *** |
|  | Age 6 * Diatom | 1.318 | 0.335 | 3.935 | 8.33e-5 | *** |
